# Comparative Analysis of Human Coronaviruses Focusing on Nucleotide Variability and Synonymous Codon Usage Pattern

**DOI:** 10.1101/2020.07.28.224386

**Authors:** Jayanta Kumar Das, Swarup Roy

**Affiliations:** Department of Pediatrics, School of Medicine, Johns Hopkins University, MD-21205, USA; Network Reconstruction & Analysis (NetRA) Lab, Department of Computer Applications, Sikkim University, Gangtok, India

**Keywords:** Nucleotide, Codon, RSCU, ENC, Amino acid, Correlation, Phylogeny, Human Coronavirus

## Abstract

Prevailing pandemic across the world due to SARSCoV-2 drawing great attention towards discovering its evolutionary origin. We perform an exploratory study to understand the variability of the whole coding region of possible proximal evolutionary neighbours of SARSCoV-2. We consider seven (07) human coronavirus strains from six different species as a candidate for our study.

First, we observe a good variability of nucleotides across candidate strains. We did not find a significant variation of GC content across the strains for codon position first and second. However, we interestingly see huge variability of GC-content in codon position 3rd (GC3), and pairwise mean GC-content (SARSCoV, MERSCoV), and (SARSCoV-2, hCoV229E) are quite closer. While observing the relative abundance of dinucleotide feature, we find a shared typical genetic pattern, i.e., high usage of GC and CT nucleotide pair at the first two positions (P12) of codons and the last two positions (P23) of codons, respectively. We also observe a low abundance of CG pair that might help in their evolution bio-process. Secondly, Considering RSCU score, we find a substantial similarity for mild class coronaviruses, i.e., hCoVOC43, hCoVHKU1, and hCoVNL63 based on their codon hit with high RSCU value (≥ 1.5), and minim number of codons hit (count-9) is observed for MERSCoV. We see seven codons ATT, ACT, TCT, CCT, GTT, GCT and GGT with high RSCU value, which are common in all seven strains. These codons are mostly from Aliphatic and Hydroxyl amino acid group. A phylogenetic tree built using RSCU feature reveals proximity among hCoVOC43 and hCoV229E (mild). Thirdly, we perform linear regression analysis among GC content in different codon position and ENC value. We observe a strong correlation (significant p-value) between GC2 and GC3 for SARSCoV-2, hCoV229E and hCoVNL63, and between GC1 and GC3 for hCoV229E, hCoVNL63, SARSCoV. We believe that our findings will help in understanding the mechanism of human coronavirus.

## 1. Introduction

Coronavirus is a large enveloped virus (family-*Coronaviridae*, subfamily-*Coronavirinae*) with non-segmented, single-stranded, and positive-sense RNA genomes [1]. Among many, six coronaviruses have been known to infect human hosts and cause respiratory diseases. The SARS (Severe Acute Respiratory Syndrome) and MERS (Middle East Respiratory Syndrome) are the most lethal coronaviruses. *SARSCoV* is the first reported in China (2002) [2, 3], which causes around two thousand deaths worldwide. *MERSCoV* is reported in Saudi Arabia (2012) and South Korea (2015) [4, 5]. In a timeline, *SARSCoV-2* is the recently reported coronavirus (2019), which provoked a large-scale epidemic, COVID-19, originated from Whung, the largest metropolitan area in China’s Hubei province. Even then the diseases they are responsible for are quite different. COVID-19 has a strong infection power due to its high dissemination rate across the worldwide. According to WHO report ^1^, more than six hundred thousand people are already dead so far (as on July 15, 2020) due to COVID-19. All these three groups of coronaviruses are highly pathogenic that resulted global outbreaks. The other three human coronaviruses are Human coronavirus OC43 (*hCoVOC43*), Human coronavirus HKU1 (*hCoVHKU1*), Human coronavirus 229E (*hCoV229E*) and Human coronavirus NL63 (*hCoVNL63*). They are categorized as mild due to its low infection and mortally rate.

From the genetic composition view point, the complete genome length of SARSCoV, SARSCoV-2, and MERSCoV is approximately 27-30 kb. The genome of SARSCoV-2 showing quite high similarity (≈ 79%) to SARSCoV and moderate similarity (≈ 50%) to MERSCoV [6]. A number of putative coding regions are available in SARSCoV-2 that encodes important genes that includes nonstructural proteins such as orf1ab, structural proteins namely spike glycoprotein (S), envelope (E), membrane (M) and nucleocapsid (N), and several accessory protein chains [7, 2, 8, 9]. The two-thirds of the genome is at the 5′ side of the sequence, encoding the nonstructural proteins, and one-third are at the 3′ side encoding four structural proteins [7]. Coronavirus proteins play diverse functional role. Nonstructural proteins can block the host innate immune response [10]. Among many structural proteins, the envelope protein promotes viral assembly and release [11]. The spike proteins compose the spikes on the viral surface, binds to host receptors [11]. However, many of these sequence variability features of structural and nonstructural proteins are yet to investigate thoroughly.

The 20 standard amino acids are genetically coded by 64 codons, including 3 stop codons for translation termination signal [12]. Therefore, a single amino acid are coded by multiple codons, which are called synonymous codons. The number of synonymous codons is varying between 1 to 6. Virus genomes are differ from each others due to frequent mutations that can prevent the PCR primers bind to target sequences [13]. Mutation plays a key role that triggered a zoonotic virus to jump from animal host to humans[14] host. Due to several other biological factors, such as pathogenicity the range of target hosts also vary diversely even in closely related strains [15, 16]. Genome-wide codon usage signature can reflect evolutionary forces [17, 17]. It has been observed that the inter and intra-species codon usage patters are varying significantly [18] in different organisms. It is therefore genetically important to study the nucleotide base composition in all the three positions of a codon, because nucleotide composition can influence the codon usage and mutational bias [19, 20]. The frequency of dinucleotide feature is also important that may affect codon usage [19]. In this context, GC content in codons may be a good indicator towards understanding expression of viral genes while interacting inside human host cells [21, 22]. Previous studies have demonstrated that usage of synonymous codon is a non-random procedure [23, 24]. The relative synonymous codon usage (RSCU) is used to standardize the codon usage of those amino acid encoded by multiple codons. The RSCU value is independent of the amino acid composition and has been used widely to estimate the codon usage bias. Several studies have been done on coronaviruses, mostly by focusing on independent genome [25, 19], different strains within the same genome [26], and host specific adaption and proximal origin of SARSCoV-2 virus [27, 28, 29]. The evolutionary mechanism of the SARSCoV, SARSCoV-2, MERSCoV, including its mutation rate, have been studied, although several crucial roles are yet to known, especially SARSCoV-2 to fight with COVID19 disease. Therefore, a comparative study on human coronaviruses might be helpful to exploit crucial factors.

The current study have focused on comprehensive and integrated way of analysing of all known seven strains of human coronaviruses. Several bias indexing parameters have used such as nucleotide composition and dinucleotide odd ratio [30], synonymous codon usage pattern [31], effective number of codon usage [32, 33], and details are discussed in the following section.

## 2. Materials and Methods

### 2.1. Data retrieval and filtering

We took all six species of known human coronaviruses, a total of seven strains as one species is divided into two different strains. Human coronaviruses can broadly be divided into two classes based on their produced symptoms in humans, either potentially severe or mild. MERSCoV, SARSCoV and SARSCoV-2 are categorized into potentially severe class. Human coronavirus OC43 (hCoVOC43), Human coronavirus HKU1 (hCoVHKU1), Human coronavirus 229E (hCoV229E) and Human coronavirus NL63 (hCoVNL63) are categorized into mild condition class. Among, MERSCoV, SARSCoV, SARSCoV-2, hCoVOC43, hCoV229E are belonging to the *Genus β*-CoV, whereas hCoV229E and hCoVNL63 are belonging to the *Genus α*-CoV.

Nucleotide sequences for each strain of length ≈ 28*kb* are collected from NCBI database during April, 2020. At first, genome wise all the partial, incomplete, and duplicate sequence are removed. Then for each complete sequence, single coding sequences are obtained by concatenating coding region (i.e., codons) of all genes (that include nonstructural polyprotein: orf1ab, structural proteins: Spike (S), Envelope (E), Membrane (M), Nucleocapsid (N) and several accessories proteins: Orf3a, Orf3b, Orf6, Orf7a, Orf7b, Orf8(a/b), Orf10) of length having ≥150 bp. The gene-wise final dataset, including the number of unique sequence count and associated coding gene, i.e., protein information are reported in Table 1, which are utilized in our subsequent analysis.

**Table 1:**
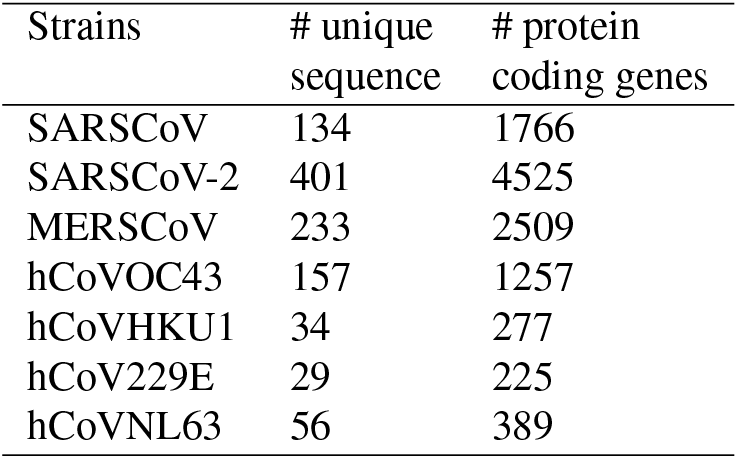
The quantitative information for number of unique sequences and involved protein coding genes for all seven strains.

### 2.2. Measuring indexes

#### Quantifying nucleotide composition

The quantity of nucleotide base composition i.e. A, T, C and G in different codon positions 1st/2nd/3rd can be calculated based on frequency measure. The base composition and GC content at first, second, and third positions of synonymously variable sense codons which can potentially vary from 0 to 1. The nucleotide composition of any nucleotide *X* at position *p* is calculated as follows:

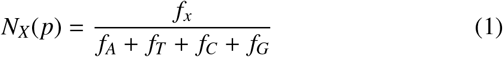

where *X* ∈ {*A*, *T*, *C*, *G*}, p∈ 1, 2, 3 and *f*_*A*_, *f*_*T*_, *f*_*C*_, *f*_*G*_ are the nucleotide frequencies at particular position *p* for A, T, C and G respectively.

Similarly, we calculate pair nucleotide composition in a particular position *p* as follows:

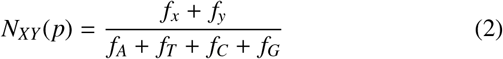

where *X*, *Y* ∈ {*A*, *T*, *C*, *G*}, p∈ 1, 2, 3 and *f*_*A*_, *f*_*T*_, *f*_*C*_, *f*_*G*_ are the nucleotide frequencies at particular position *p* for A, T, C and G respectively.

#### Relative dinucleotide abundance

Dinucleotide frequency is often used to determine favourable or unfavourable dinucleotide pairs. The variation in the frequency of dinucleotide pairs may affect codon usage. The total possible dinucleotide combinations are 16. The patterns of dinucleotide frequency indicate both selectional and mutational pressures. Relative Dinucleotide Abundance frequency is calculated using the following formula:

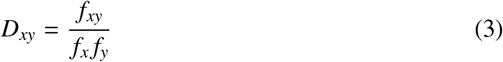

where *f*_*x*_ and *f*_*y*_ represent the individual frequency of nucleotides *x* and *y* respectively, and *f*_*x*_*y* is the frequency of dinucleotides (xy) in the same sequence. We calculate relative dinucleotide frequency for the codon positions first and second as well as second and third. The ratio of the observed to expected dinucleotide frequency is known as the odds ratio. The value of the odds ratio indicates underrepresented, if the value is below 0.78, and overrepresented, if the value is above 1.25 [30].

#### Relative synonymous codon usage (RSCU) pattern

Relative synonymous codon usage (RSCU) is the observed number of occurrences of codons divided by that expected if the usage of synonymous codons uniformly [31] (Equation 4). The RSCU is used to standardize the codon usage of those amino acid encoded by multiple codons. The RSCU value is independent of the amino acid composition and has been used widely to estimate the codon usage bias. The RSCU value greater than 1.0 is considered to be a positive codon usage bias, and the RSCU value is less than 1.0 is considered to be a negative codon usage bias. Thus, a higher RSCU value means that the codon is used more frequently.

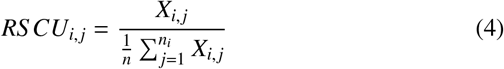

where *X*_*i*_ is the number of occurrences of the *jth* codon for the *ith* amino acid, which is encoded by *n*_*i*_ synonymous codons.

#### Effective number of codon (*ENC*) usage

The effective number of codon (*ENC*) usage can be obtain by Equation 5 [32, 33].

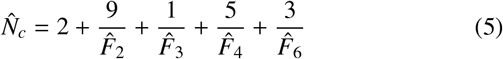

where *F*_*i*_ denotes the average homozygosity for the class with *i* synonymous codons. The *ENC* value ranges from 20 to 61. An *ENC* of 20 represents extreme bias as only one codon is used for each amino acid, and a value of 61 suggests that there is no bias. In contrast to the RSCU value, a higher *ENC* value correlates to a weaker codon usage bias. An alternate approach for calculating the *ENC* is based on the GC3 content given in Equation 6.

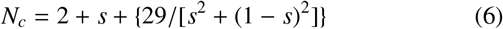

## 3. Results and Discussion

### 3.1. Variability of nucleotide composition

The variability of nucleotide composition is shown by box plot in Figure 1. It is noted that strain wise nucleotide composition significantly differs, where base composition for the MERSCoV genome is highly diverse for all nucleotide bases. Here, we focus a few high or low content of nucleotide composition for different strains. We highlight each base of nucleotide content. We observe that the content of A is high for SARSCoV-2, and is low for MERSCoV and hCoVNL63; the content of T is high for hCoVHKU1 and hCoVNL63, and is low for SARSCoV; the content of C is high for SARSCoV and MERSCoV, and is low hCoVHKU1; the content of G is high for hCoVOC43 and hCoV229E, and is low for hCoVHKU1. General trends of CDSs of all coronavirus strains are shown to be rich in A and T (≈ 58-67%-), in comparison to G and C nucleotide., with the maximum is for hCoVHKU1 and minimum is for MERSCoV. If A is high, then T is comparatively low and vice versa. This characteristic is similar to that of Nipah virus [19].

**Figure 1:**
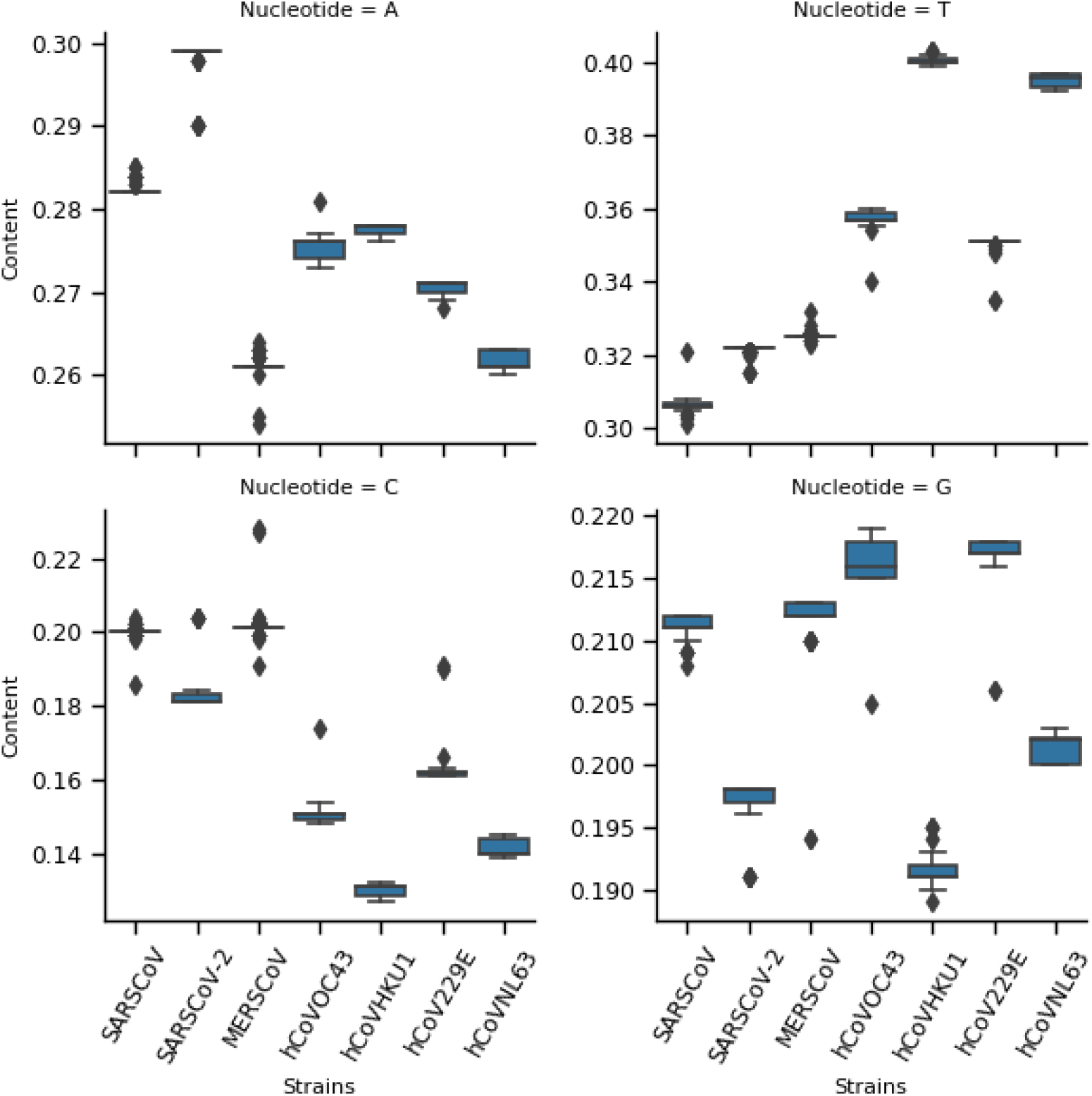
The box showing the distribution of A/T/C/G content for seven strains of human coronaviruses.

### 3.2. Dinucleotide variability and usage pattern

Dual-nucleotide (DN) composition and its variability are also essential as they represent the possible bonding nature and abundance of two consecutive nucleotides over the sequences. The occurrence of dinucleotide is important to study that directly play a role by impact on Codon usage bias, mutational bias and also the influence of selective pressure and compositional constraints. In RNA viruses, the relative abundance of dinucleotide has shown to affect codon usage [20]. We focus on both the first two codon position (P12) and the last two codon position (P23) for a quantitative measure of consecutive nucleotide pair, and the observed frequency distribution of nucleotide pair is shown in Figure 2. In the present study, high abundance nucleotide pairs are AA, GA, GT and TT for P12 as compared to high abundance nucleotide pair AT, CT, GT and TT for P23. Among these dinucleotides, GA and CT belong to purine and pyrimidine group, respectively. We also observe less variability of nucleotide pair across the strains for P23 compared to P12 (Figure 2). Overall, this indicates a distinct pattern of nucleotide pairs in two consecutive codon positions.

**Figure 2:**
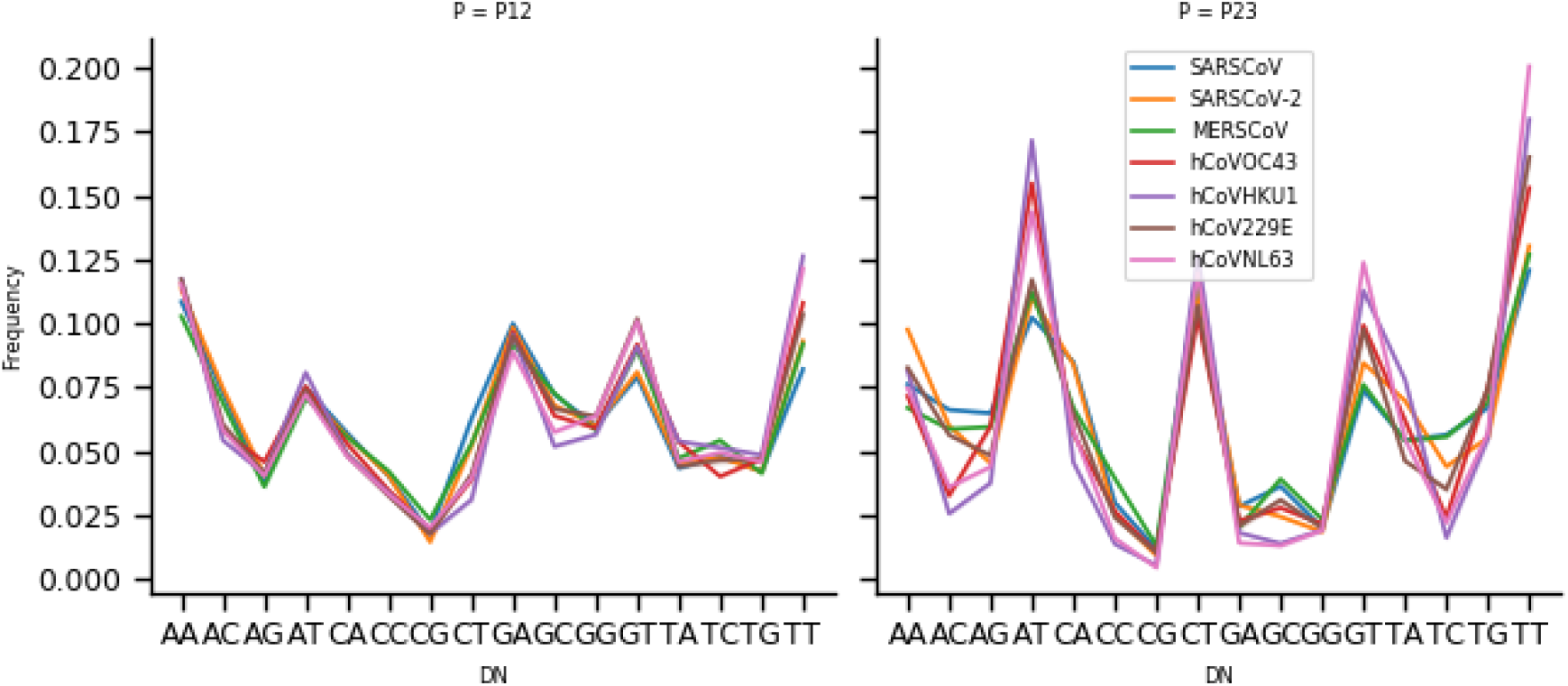
Distribution of average di-nucleotide variability for seven strains of human coronaviruses.

We measure the relative abundance of dinucleotide content (odd ratio) as shown in Table 2. The top two high and low usage dinucleotide pair is provided. Quantitatively, we observe a similar pattern for the highest and lowest usage nucleotide pairs, which are GC and CG, respectively, across the seven strains for P12. However, we find different structures for the second-highest and lowest usage nucleotide pair. We also observe a distinct arrangement of high and low usage nucleotide pairs for P23 across the strains and that is also different from the position P12 as stated above.

**Table 2:**
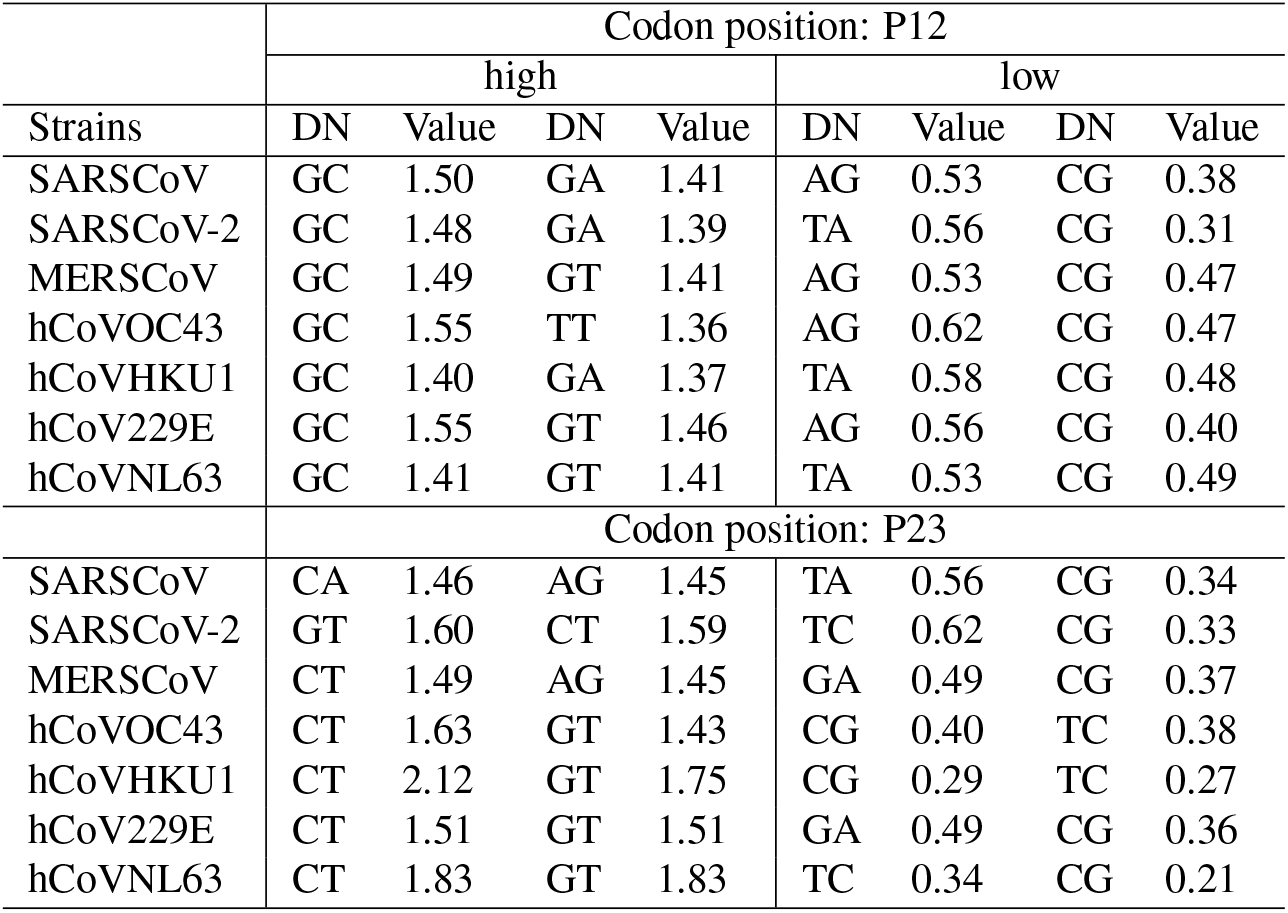
The top two high and low usage dinucleotide odd ratio value for the both combination of codon positions: P12 and P23.

### 3.3. GC-content usage pattern

GC-content variability by the mean and standard deviation in three different positions (GC1, GC2, GC3) for seven strains of coronavirus is shown in Table 3, which is calculated using Eq. 2. We interestingly observe that GC3 shows high variability among seven strains, although the GC-content for GC1 and GC2 is much greater than that of GC3. Pairwise, mean GC-content (SARSCoV, MSERCoV) and (SARSCoV-2, hCoV229E) are closer. We then look the position-wise GC-content distribution (Figure 3). It is observed that GC-content in three codon positions is well balanced for hCoVOC43 and SARSCoV-2 (mean difference ≈ 0.9). Whereas, hCoVHKU1 and hCoVNL63 show closer GC-content between the first and second positions of codon (mean difference ¡ 0.9), and SARSCoV, MERSCoV and hCoV229E show closer GC-content between the second and third positions of codon (mean difference < 0.9).

**Table 3:**
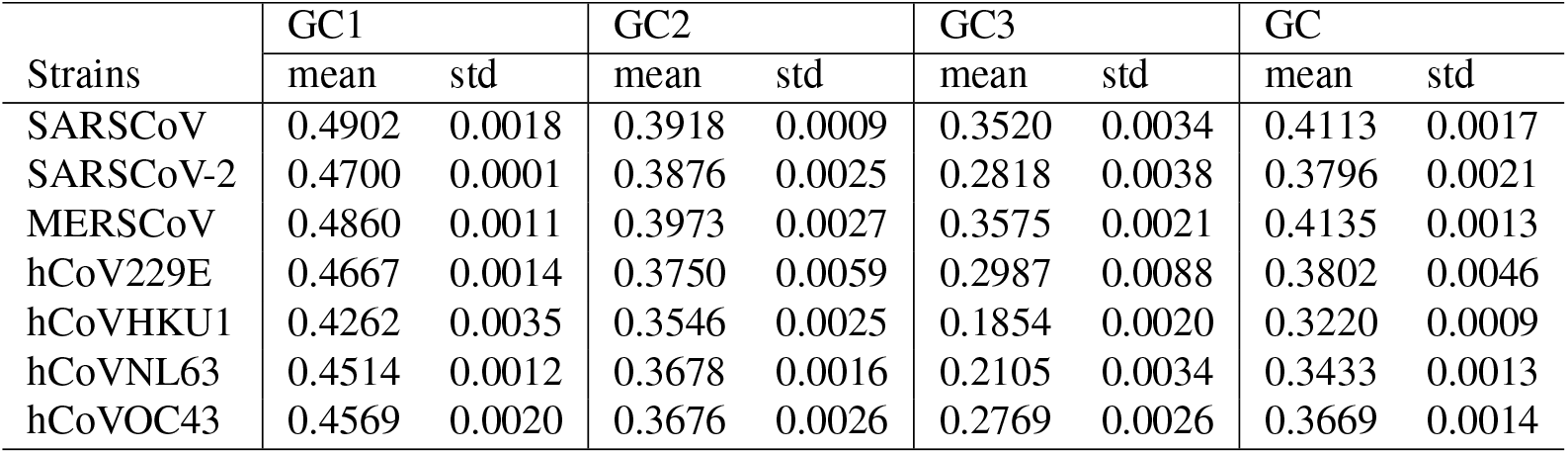
GC content variability- mean and std. for each position of GC content for seven strains of coronaviruses.

**Figure 3:**
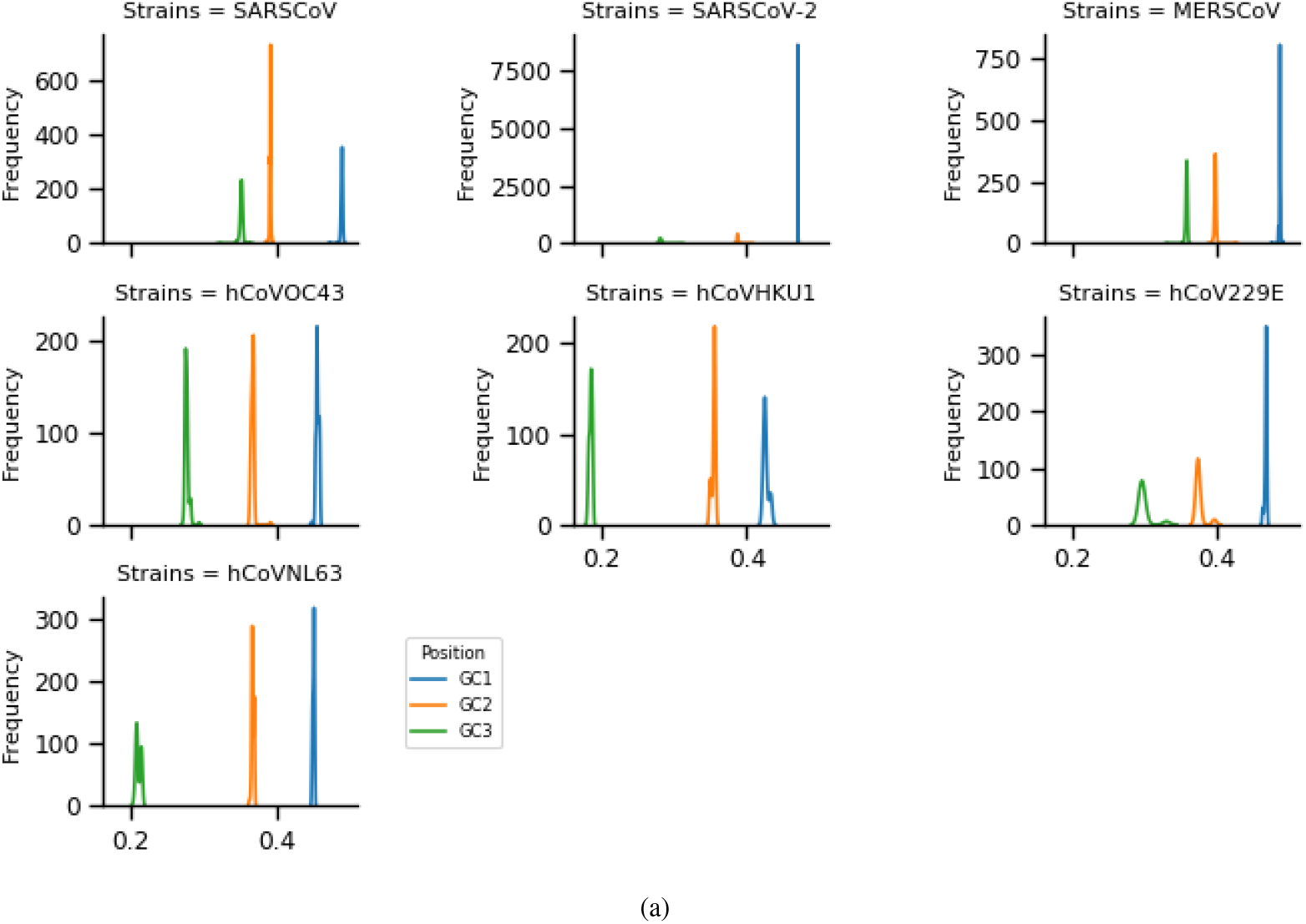
Distribution in position wise (first-GC1, second-GC2, third-GC3, and first and second combined-GC12) GC-content composition for seven strains of human coronaviruses.

### 3.4. Synonymous codon usage pattern and phylogenetic clustering

We obtain the RSCU values of 59-non trivial codons (Eq. 4). Strains-wise, RSCU value for each codon is shown in Table 4. RSCU value with a high score (≥ 1.5) is highlighted for all seven strains. We observe the maximum number of codons hit (count-18) whose RSCU value is high in three strains of mild class coronaviruses hCoVOC43, hCoVHKU1, and hCoVNL63 and minim number of codons hit (count-9) is observed for MERSCoV. Among the highlighted, we observe only seven codons ATT, ACT, TCT, CCT, GTT, GCT and GGT, which are common for all seven strains with high RSCU value (≥ 1.5). We observe a few codons, which are highly utilized (based on RSCU value), can be categorized into a different group of amino acids [34] (Figure 4). Such as five from Aliphatic group (A-GCT, G-GGT, I-ATT, L-TTA, V-GTT), two from hydroxyl containing group (T-TCT, S-ACT), one from Basic amino acid group (R-AGA, R- CGT) and one from Cyclic group (P-CCT). Out of those seven highly common usage codons, four are from Aliphatic group (I-ATT, V-GTT, A-GCT, G-GGT), one from Cyclic group (P-CCT) and one from Sulfur-containing group (T-ACT) [34].

**Table 4:** Relative synonymous codon usage (RSCU) analysis of the various codons of all coding sequences combined in seven strains of coronaviruses. AA-Amino acid, Highlighted cells are those with RSCU value> 1.5.

| A A | Codon | SARSCoV | SARSCoV-2 | MERSCoV | hCoVOC43 | hCoVHKU1 | hCoV229E | hCoVNL63 |
| --- | --- | --- | --- | --- | --- | --- | --- | --- |
| K | AAA | 1.042 | 1.295 | 1.004 | 1.041 | 1.36 | 1.133 | 1.203 |
| N | AAT | 1.236 | 1.36 | 1.395 | 1.659 | 1.742 | 1.379 | 1.611 |
| N | AAC | 0.764 | 0.64 | 0.605 | 0.341 | 0.258 | 0.621 | 0.389 |
| K | AAG | 0.958 | 0.705 | 0.996 | 0.959 | 0.64 | 0.867 | 0.797 |
| I | ATA | 0.625 | 0.921 | 0.711 | 0.97 | 0.892 | 0.748 | 0.676 |
| I | ATT | 1.71 | 1.525 | 1.718 | 1.756 | 1.94 | 1.878 | 2.128 |
| I | ATC | 0.665 | 0.554 | 0.572 | 0.274 | 0.167 | 0.373 | 0.196 |
| T | ACA | 1.571 | 1.641 | 1.185 | 1.27 | 1.051 | 1.472 | 1.204 |
| T | ACT | 1.664 | 1.797 | 1.952 | 1.976 | 2.616 | 1.904 | 2.376 |
| T | ACC | 0.583 | 0.38 | 0.686 | 0.54 | 0.249 | 0.432 | 0.325 |
| T | ACG | 0.182 | 0.183 | 0.177 | 0.214 | 0.083 | 0.192 | 0.095 |
| R | AGA | 2.082 | 2.642 | 1.347 | 1.84 | 2.046 | 2.224 | 1.427 |
| S | AGT | 1.172 | 1.463 | 1.336 | 2.075 | 1.916 | 1.621 | 1.947 |
| S | AGC | 0.523 | 0.335 | 0.437 | 0.575 | 0.188 | 0.504 | 0.236 |
| R | AGG | 0.927 | 0.831 | 0.842 | 0.663 | 0.56 | 0.628 | 0.823 |
| Y | TAT | 1.122 | 1.205 | 1.27 | 1.669 | 1.778 | 1.378 | 1.629 |
| Y | TAC | 0.878 | 0.795 | 0.73 | 0.331 | 0.222 | 0.622 | 0.371 |
| L | TTA | 1.042 | 1.63 | 1.216 | 1.506 | 2.397 | 1.127 | 1.701 |
| F | TTT | 1.231 | 1.43 | 1.284 | 1.752 | 1.847 | 1.63 | 1.802 |
| F | TTC | 0.769 | 0.57 | 0.716 | 0.248 | 0.153 | 0.37 | 0.198 |
| L | TTG | 1.084 | 1.099 | 1.433 | 1.928 | 1.637 | 2.064 | 1.838 |
| S | TCA | 1.694 | 1.666 | 1.217 | 0.858 | 0.869 | 1.102 | 1.136 |
| S | TCT | 1.959 | 1.971 | 2.11 | 1.84 | 2.697 | 2.146 | 2.343 |
| S | TCC | 0.423 | 0.458 | 0.712 | 0.463 | 0.234 | 0.479 | 0.246 |
| S | TCG | 0.229 | 0.108 | 0.189 | 0.189 | 0.097 | 0.149 | 0.091 |
| C | TGT | 1.265 | 1.575 | 1.194 | 1.534 | 1.83 | 1.462 | 1.821 |
| C | TGC | 0.735 | 0.425 | 0.806 | 0.466 | 0.17 | 0.538 | 0.179 |
| Q | CAA | 1.174 | 1.368 | 1.141 | 1.072 | 1.363 | 1.32 | 1.344 |
| H | CAT | 1.29 | 1.396 | 1.32 | 1.581 | 1.764 | 1.457 | 1.612 |
| H | CAC | 0.71 | 0.604 | 0.68 | 0.419 | 0.236 | 0.543 | 0.388 |
| Q | CAG | 0.826 | 0.632 | 0.859 | 0.928 | 0.637 | 0.68 | 0.656 |
| L | CTA | 0.645 | 0.647 | 0.459 | 0.414 | 0.281 | 0.355 | 0.254 |
| L | CTT | 1.782 | 1.763 | 1.698 | 1.433 | 1.324 | 1.764 | 1.876 |
| L | CTC | 0.84 | 0.577 | 0.702 | 0.297 | 0.161 | 0.303 | 0.208 |
| L | CTG | 0.606 | 0.285 | 0.492 | 0.423 | 0.199 | 0.387 | 0.123 |
| P | CCA | 1.687 | 1.587 | 1.219 | 1.214 | 0.943 | 1.326 | 1.229 |
| P | CCT | 1.744 | 1.973 | 1.945 | 2.069 | 2.605 | 2.057 | 2.506 |
| P | CCC | 0.406 | 0.29 | 0.646 | 0.503 | 0.303 | 0.441 | 0.174 |
| P | CCG | 0.162 | 0.149 | 0.19 | 0.214 | 0.149 | 0.176 | 0.091 |
| R | CGA | 0.428 | 0.3 | 0.449 | 0.455 | 0.342 | 0.244 | 0.272 |
| R | CGT | 1.739 | 1.469 | 1.813 | 1.977 | 2.354 | 2.14 | 2.894 |
| R | CGC | 0.727 | 0.579 | 1.114 | 0.728 | 0.462 | 0.635 | 0.485 |
| R | CGG | 0.097 | 0.18 | 0.435 | 0.338 | 0.236 | 0.129 | 0.1 |
| E | GAA | 1.048 | 1.448 | 1.052 | 1.189 | 1.395 | 1.407 | 1.284 |
| D | GAT | 1.241 | 1.275 | 1.277 | 1.657 | 1.704 | 1.277 | 1.567 |
| D | GAC | 0.759 | 0.725 | 0.723 | 0.343 | 0.296 | 0.723 | 0.433 |
| E | GAG | 0.952 | 0.552 | 0.948 | 0.811 | 0.605 | 0.593 | 0.716 |
| V | GTA | 0.832 | 0.888 | 0.724 | 0.682 | 0.79 | 0.445 | 0.445 |
| V | GTT | 1.718 | 1.984 | 1.778 | 2.216 | 2.726 | 2.338 | 2.906 |
| V | GTC | 0.668 | 0.543 | 0.762 | 0.318 | 0.232 | 0.494 | 0.332 |
| V | GTG | 0.782 | 0.586 | 0.736 | 0.785 | 0.252 | 0.723 | 0.317 |
| A | GCA | 1.129 | 1.09 | 0.989 | 1.106 | 0.917 | 1.142 | 1.078 |
| A | GCT | 2.069 | 2.175 | 2.069 | 2.159 | 2.622 | 2.167 | 2.395 |
| A | GCC | 0.574 | 0.578 | 0.631 | 0.538 | 0.347 | 0.467 | 0.448 |
| A | GCG | 0.227 | 0.157 | 0.312 | 0.197 | 0.114 | 0.225 | 0.079 |
| G | GGA | 0.867 | 0.791 | 0.644 | 0.673 | 0.418 | 0.409 | 0.31 |
| G | GGT | 2.015 | 2.403 | 2.063 | 2.492 | 3.028 | 2.745 | 3.33 |
| G | GGC | 0.943 | 0.693 | 1 | 0.593 | 0.405 | 0.754 | 0.272 |
| G | GGG | 0.175 | 0.112 | 0.293 | 0.242 | 0.149 |  |  |

**Figure 4:**
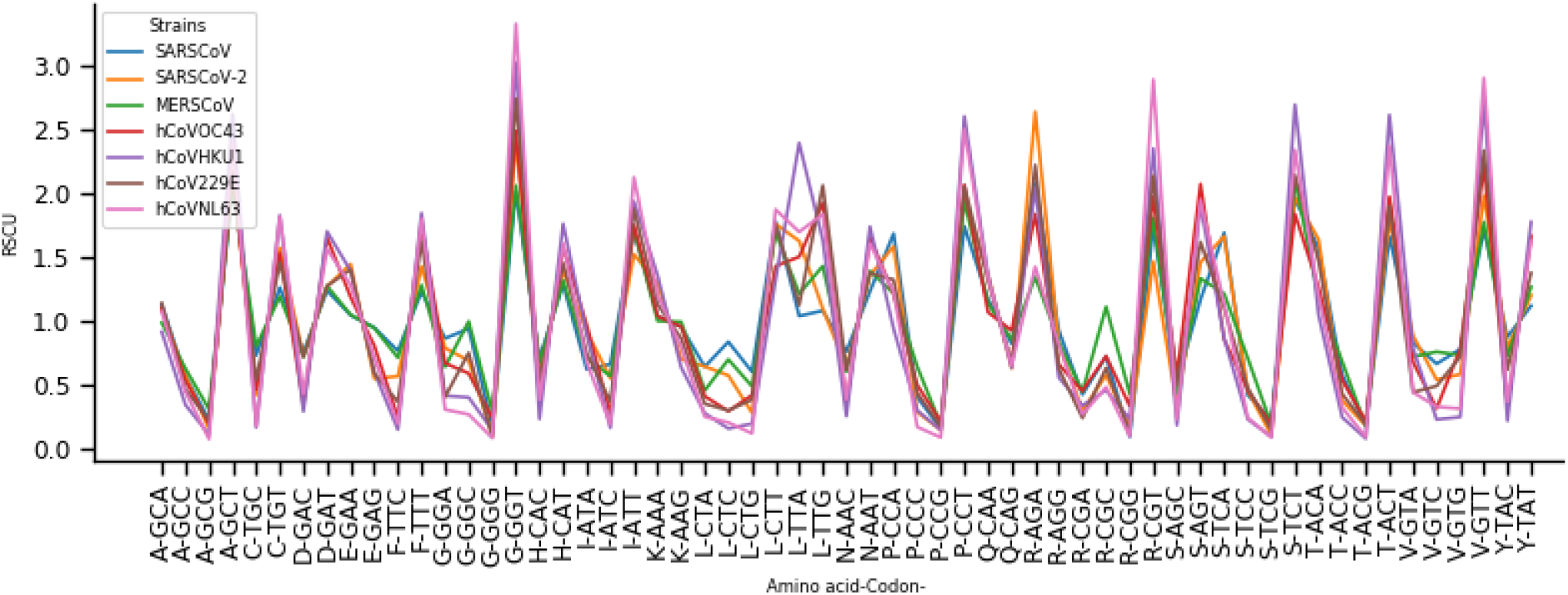
Distribution of RSCU value shown by amino acid with synonymous codons.

To understand the evolutionary mechanism, we consider RSCU values as a feature vector. We obtain average RSCU value for each codon within the multiple sequences of the same strain and obtain a strainwise 59-dimensional feature vector. We perform hierarchical clustering (dendogram) with the ‘average’ linkage method, and the obtain graph is shown in Figure5. It is observed that the severe and mild class coronaviruses are in different clades. However, hCoVOC43 and hCoV229E are very close to those of the severe class coronaviruses. This is an indication of similar codon usage patterns that might be a good indicator of findings difference.

**Figure 5:**
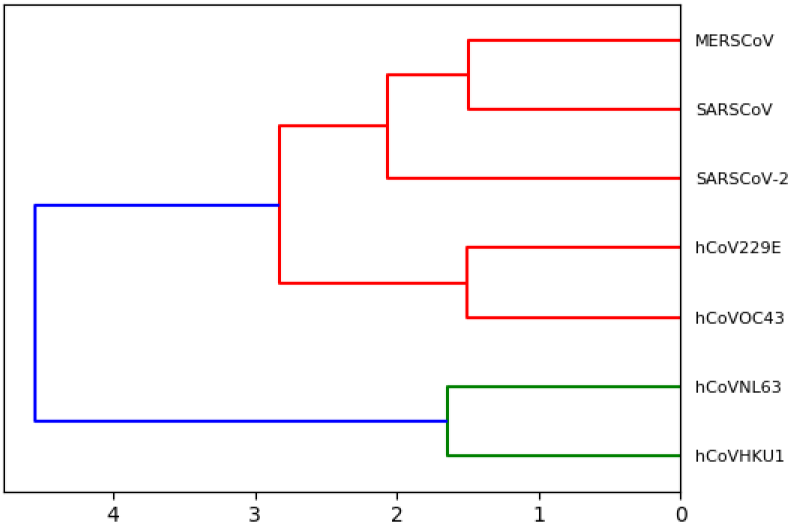
Hierarchical clustering of all seven strains of human coronaviruses using RSCU vector.

### 3.5. Performing correlation analysis using different indexing measure

To account for the factor that might influence the low or high codon usage bias, we calculate ENC value for all seven strains. We observe mean ENC value is ranging from 36.40 (for hCoVHKU1) to 49.8 (for MERSCoV). An ENC value greater than 45 is considered as a lower codon usage bias. We observe mean ENC value for MERSCoV and SARSCoV is relatively higher than that of SARSCoV-2 and other coronaviruses. However, ENC values for mild class coronavirus hCoVOC43 (ENC:43.794) and hCoV229E (ENC:43.1) are much higher than that of ENC values for hCoV229E(ENC:36.4) and hCoVNL63 (ENC:37.32). The previous studies on SARSCoV and MERSCoV coronaviruses have been confirmed these findings as reported in [35, 26]. Several other studies on different viruses have shown the usage of low codon bias for influenza A virus [36], Classical swine fever virus[37] and high codon bias for Hepatitis A virus [38]. Further, that might influence the low codon usage bias, and we analyzed the relationship between the ENC value and the GC-content in the third site of codons (GC3) in all seven strains of coronavirus genomes. In Figure 6, the solid line represents the expected curve. We find all the proportion of points lying near to the solid line on the left region i.e. observed value was smaller than the expected value. These findings suggest that mutational bias might have a role in determining the codon usage variation. Therefore, the codon usage bias for severe class coronaviruses is relatively low, hence low codon usage bias. This study has also confirmed the additional role of dinucleotide abundance for the evolution of severe class coronaviruses.

**Figure 6:**
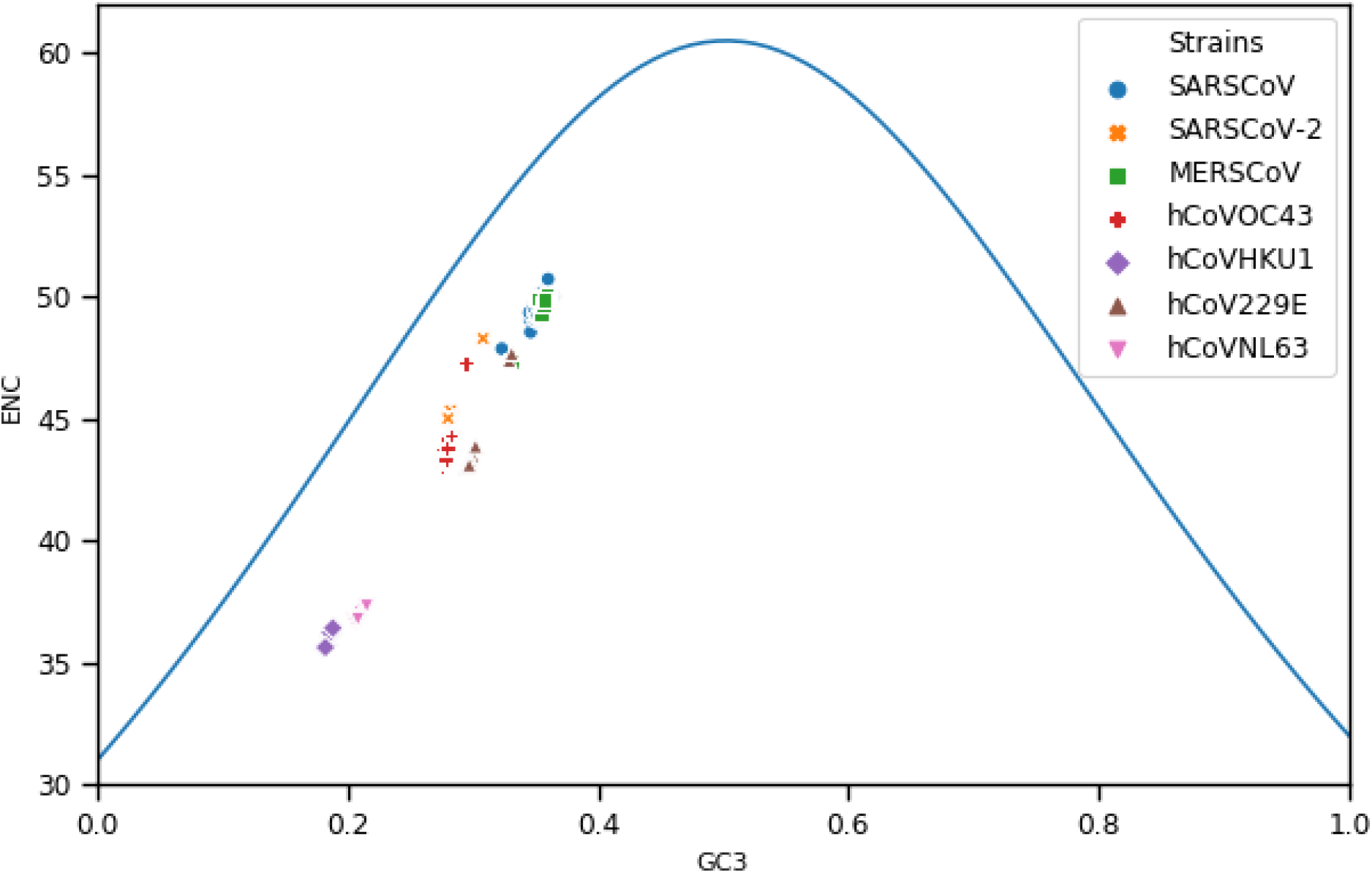
The plots of ENC values against GC3s values for all seven strains shown by different colors and style.

In synonymous codons, only the last nucleotide is different (except two codons of each Arginine and Serine amino acid), and the amino acid remained unchanged. So, the changes of the nucleotide at the third position, it is indicative of a mutational force only. Only when nucleotide change at the 2nd and third position of codons, it leads to selection force. Towards understanding this fact, we next perform a linear regression analysis for GC-content at third i.e., GC3 with the GC-content at first position (GC1) and the second position (GC2), and taking both i.e., GC12. First, we show the distribution of average GC-content for seven strains of human coronaviruses (Figure 7(a)), From the figure, we observe various overlapping region between SARSCoV-2 and hCoV229E, and SARSCoV and MERSCoV, identifying similar GC-content pattern. It can also observe that position-wise distribution can varying among strains (Figure 7(b)). GC content at the third position of codons (GC3) is a putative indicator of the extent of base composition. Therefore, we perform a linear regression analysis, GC3 with GC12, GC1 and GC2. The neutrality plot analysis of GC3 against GC1/GC2/GC12 for all seven strains of coronaviruses is shown in Figure 8. The solid line in the figure represents the regression line. The detail of the regression line with significant statistical p-value, and *R*^2^ value is shown in Table 5. The slope of the regression line suggests that the relative neutrality (mutation pressure) for GC1/GC2/GC12 and that the relative constraint on the GC3 (natural selection). It is observed that there a strong correlation between GC2 and GC3 both for SARSCoV-2 (*R*^2^ = 0.96,*p* < 0.001) and hCoV229E (*R*^2^ = 0.97,*p* < 0.001) and a moderate correlation for hCoVNL63 (*R*^2^ = 0.81,*p* < 0.001). In comparing GC3 and GC1, we observe comparatively low correlation, hCoV229E (*R*^2^ = 0.70,*p* < 0.001) hCoVNL63 (*R*^2^ = 0.48,*p* < 0.001) and SARSCoV (*R*^2^ = 0.37,*p* < 0.001). If there is a correlation between the GC12 and GC3 that is codon bias is present at all the codon positions, and this is likely to be due to mutational forces that influencing codon bias [39]. When comparing GC3 with GC12, we observe strong correlation for SARSCoV-2 and hCoV229E (Table 5). Above mentioned highlighted points, only for hCoV229E, we observe negative correlation (GC3 Vs. GC1).

**Table 5:** Linear regression computed by comparing GC3 with GC1, GC2 and GC12 for all seven strains of coronaviruses. Calculated correlation (*R*^2^) and *p*-value are also shown.

| Strains | GC3 Vs. | Regression line | $R^2$ | p-value |
| --- | --- | --- | --- | --- |
| SARSCoV | GC1 | $y = 0.33x + 0.38$ | 0.3768 | 0.0 |
| | GC2 | $y = 0.03x + 0.38$ | 0.0141 | 0.1713 |
| | GC12 | $y = 0.18x + 0.38$ | 0.2521 | 0.0 |
| SARSCoV-2 | GC1 | $y = 0.00x + 0.47$ | 0.0005 | 0.6602 |
| | GC2 | $y = 0.65x + 0.20$ | 0.9659 | 0.0 |
| | GC12 | $y = 0.36x + 0.33$ | 0.9695 | 0.0 |
| MERSCoV | GC1 | $y = 0.12x + 0.44$ | 0.0534 | 0.0004 |
| | GC2 | $y = -0.19x + 0.46$ | 0.022 | 0.0236 |
| | GC12 | $y = -0.02x + 0.45$ | 0.0005 | 0.7275 |
| hCoVOC43 | GC1 | $y = -0.08x + 0.48$ | 0.0098 | 0.2173 |
| | GC2 | $y = 0.30x + 0.28$ | 0.09 | 0.0001 |
| | GC12 | $y = 0.10x + 0.39$ | 0.0502 | 0.0048 |
| hCoVHKU1 | GC1 | $y = -0.82x + 0.58$ | 0.2321 | 0.0039 |
| | GC2 | $y = 0.72x + 0.22$ | 0.3552 | 0.0002 |
| | GC12 | $y = -0.02x + 0.39$ | 0.0022 | 0.7941 |
| hCoV229E | GC1 | $y = -0.14x + 0.51$ | 0.7023 | 0.0 |
| | GC2 | $y = 0.66x + 0.18$ | 0.9799 | 0.0 |
| | GC12 | $y = 0.26x + 0.34$ | 0.9311 | 0.0 |
| hCoVNL63 | GC1 | $y = -0.25x + 0.50$ | 0.4835 | 0.0 |
| | GC2 | $y = 0.43x + 0.28$ | 0.8114 | 0.0 |
| | GC12 | $y = 0.11x + 0.39$ | 0.2866 | 0.0 |

**Figure 7:**
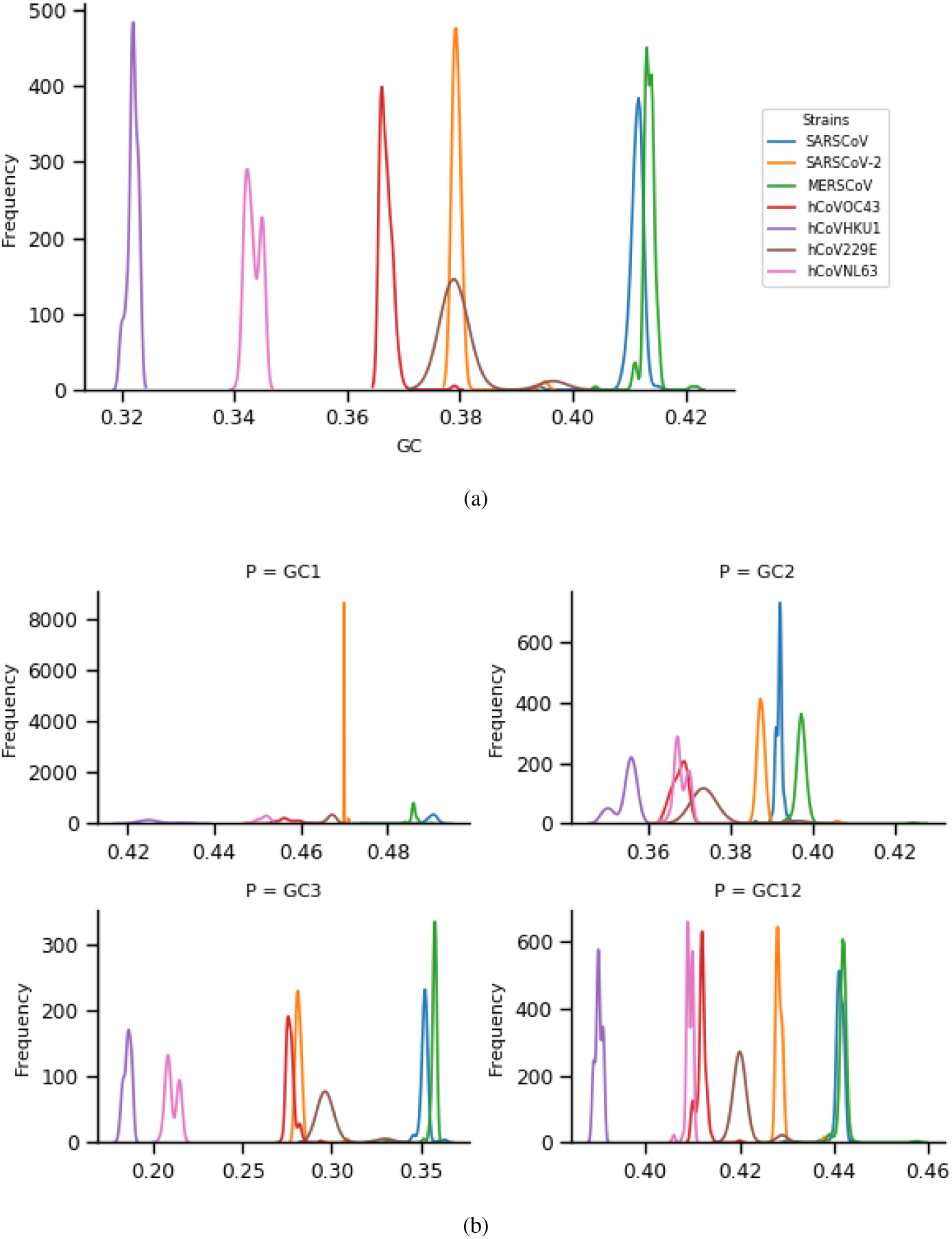
(a) Distribution of GC-content composition for seven strains of human coronaviruses. (b) Position wise distribution of GC-content (first-GC1,second-GC2,third-GC3 and first and second in combined-GC12) in seven strains of human coronaviruses.

**Figure 8:**
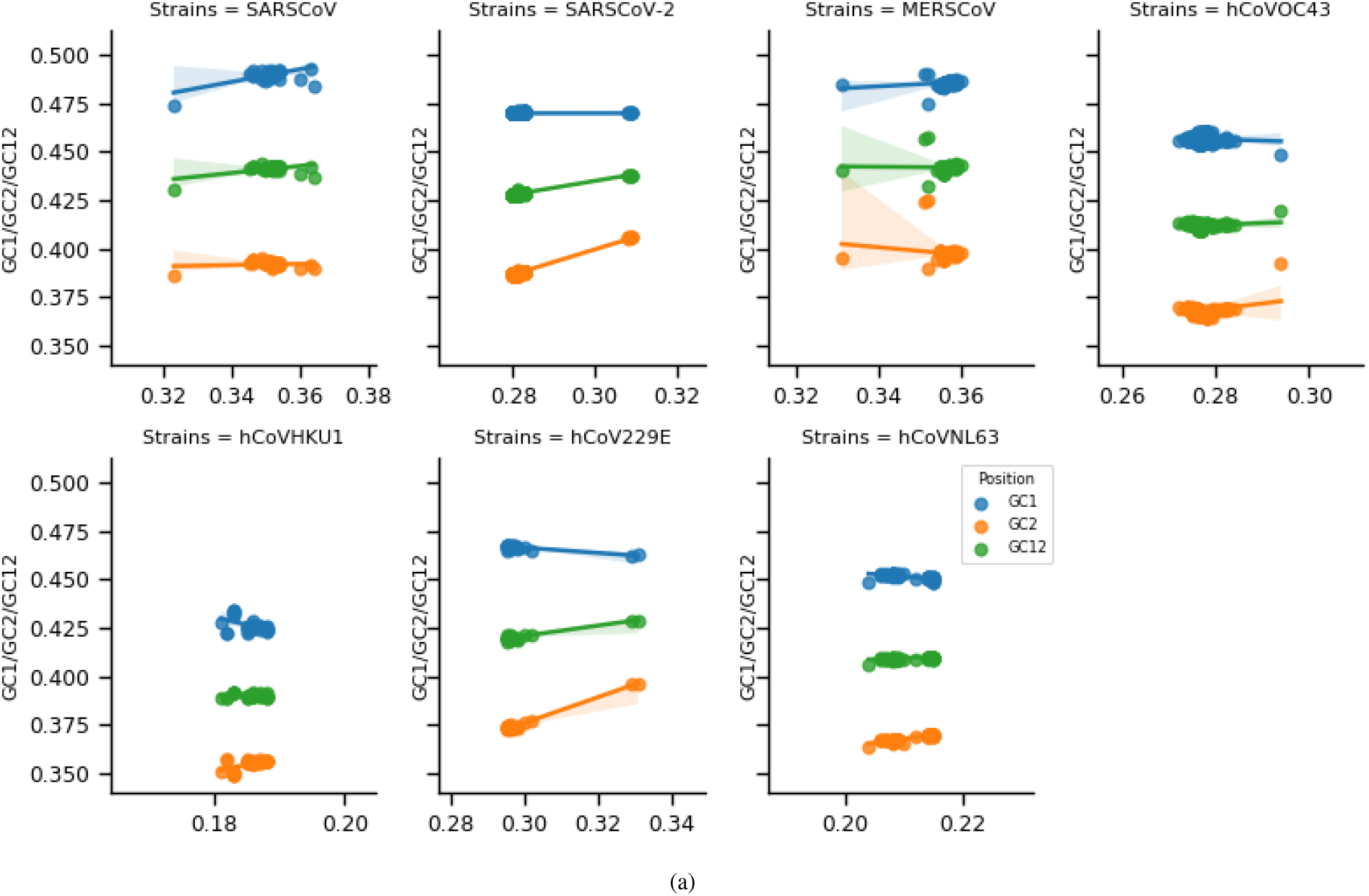
The neutral analysis of GC3 against GC1/GC2/GC12 for all seven strains of coronaviruses. The solid line represents the regression line.

## 4. Conclusion

In this work, we performed an extensive analytical and exploratory study on the genome sequence of seven human coronavirus species. We reported significant features for all candidate coronaviruses that are common and distinct. We observed that overall GC-content in codon position 3rd (GC3) is playing a crucial role in differentiating strains. We further showed that the high usage of GC (Strong group) dinucleotide in the first two positions of codons and CT (Week group) dinucleotide in the last two locations (P23) of codons. In terms of the synonymous codon usage pattern, we reported a high degree of similarity among mild class human coronaviruses. Among severe class, a shallow usage pattern observed for MERSCoV. These codons mostly belong to Aliphatic and Hydroxyl amino acid group. A phylogenetic tree built using RSCU feature reveals proximity among hCoVOC43 and hCoV229E (mild). We observed a strong correlation (significant p-value) between GC2 and GC3 for SARSCoV-2, hCoV229E and hCoVNL63, and between GC1 and GC3 for hCoV229E, hCoVNL63, SARSCoV. We believe that our findings will help in understanding the mechanism of the mutation in the host and viral genes and its consequence across the human coronavirus.

## Footnotes

1 https://www.worldometers.info/coronavirus/

